# Interrogating conserved elements of diseases using Boolean combinations of orthologous phenotypes

**DOI:** 10.1101/017947

**Authors:** John O. Woods, Matthew Z. Tien, Edward M. Marcotte

## Abstract

Conserved genetic programs often predate the homologous structures and phenotypes to which they give rise; eyes, for example, have evolved several dozen times, but their development seems to involve a common set of conserved genes. Recently, the concept of orthologous phenotypes (or phenologs) offered a quantitative way to describe this property. Phenologs are phenotypes or diseases from separate species who share an unexpectedly large set of their associated gene orthologs. It has been shown that the phenotype pairs which make up a phenolog are mutually predictive in terms of the genes involved. Recently, we demonstrated the ranking of gene–phenotype association predictions using multiple phenologs from an array of species. In this work, we demonstrate a computational method which provides a more targeted view of the conserved pathways which give rise to diseases. Our approach involves the generation of synthetic pseudophenotypes made up of Boolean combinations (union, intersection, and difference) of the gene sets for phenotypes from our database. We search for diseases that overlap significantly with these Boolean phenotypes, and find a number of highly predictive combinations. While set unions produce less specific predictions (as expected), intersection and difference-based combinations appear to offer insights into extremely specific aspects of target diseases. For example, breast cancer is predicted by zebrafish methylmercury response minus metal ion response, with predictions *MT-COI, JUN, SOD2, GADD45B*, and *BAX* all involved in the pro-apoptotic response to reactive oxygen species, thought to be a key player in cancer. We also demonstrate predictions from *Arabidopsis* Boolean phenotypes for increased brown adipose tissue in mouse (salt stress response’s intersection with sucrose stimulus response); and for human myopathy (red light response minus water deprivation response). We demonstrate the ranking of predictions for human holoprosencephaly from the set intersections between each pair of a variety of closely-related zebrafish phenotypes. Our results suggest that Boolean phenolog combinations may provide a more informed insight into the conserved pathways underlying diseases than either regular phenologs or the naïve Bayes approach.

## 1 Background

Limbs are an example of homologous structures — existing in multiple species — which appear to have evolved independently, but share underlying sets of conserved genes [1,2]. This concept, known as deep homology [1], explains the remarkable convergent evolution of eyes several dozen times [3]; the underlying genetic programs responsible for eye development must predate eyes [1]. It follows that such deeply conserved genetic processes played some selectively advantageous role in the most recent common ancestor and produced a measurable phenotype.

Phenologs are an extension of the homology of individual genes to sets of genes affiliated with specific structures, phenotypes, or diseases. McGary *et al.* defined phenologs as orthologous phenotypes — phenotypes from separate species which share an unexpectedly large number of associated genes (as determined by gene orthology). Furthermore, phenologs are mutually predictive; genes involved in one phenotype, but not known to play a role in a second, are predicted for the second, and vice versa [4]. Woods *et al.* further demonstrated that gene–phenotype association predictions using phenologs can be improved by integrating information from multiple phenologs across several species [5]. If cases of structural homology have a non-structural common ancestor, phenologs fill in the theoretical gap for determining what that common ancestor might have been.

Using phenologs, we previously demonstrated the prediction and experimental validation of vertebrate neural crest development genes such as *SEC23IP* on the basis of the plant phenotype *negative gravitropism defective* [4]. Subsequently, we provided literature validation for genes we predicted for epilepsy and atrial fibrillation from a *k* nearest neighbors search and Bayesian integration of orthologous phenotypes [5]. Whereas the *k* nearest neighbors approach yielded a broad look at the orthologous processes which give rise to diseases and phenotypes, we present in this manuscript a more focused look. Both the phenolog and deep homology hypotheses are based on a modularity postulate: proteins and genes function together in pathways or functional modules which may be composed narrowly or broadly (and thus are divisible into more narrowly-defined functionalities). Notably, Bowers *et al.* identified triplets of proteins whose phylogenetic profiles — concurrent presence, absence, or combinations of the two — obey higher order relationships than have presently been explored using phenologs [6]. We were interested in determining whether these logical relationships hold true in phenotypic space as well as functional module space.

In this work, we describe the use of Boolean combinations of phenotypes to generate pseudo-phenotypes representing hypothetical functional modules, which are often more faithfully orthologous to the query diseases than were the original components. We demonstrate the prediction of genes associated with oxidative stress-related apoptosis in breast cancer from zebrafish, and increased brown adipose tissue as well as myopathy from plants. We also provide the source code and datasets.

## 2 Methods

### 2.1 Matrix framework

We utilized NMatrix, part of the SciRuby Project, for representation of sparse matrices in the Ruby language; the storage type is known as ‘new’ Yale [7] (described more thoroughly in [8]), which stores only the diagonal and non-zero elements of each row.

As such, phenotypes may be represented as matrix rows and genes as columns; as in [5], cells containing 1 have an observed association between a gene and phenotype, and cells with 0 indicate no observation (as opposed to observed negative association, which is not included in our model). One gene–phenotype matrix is used to represent each species.

As in [5], INPARANOID orthogroups may be utilized to translate individual species-specific gene–phenotype matrices into sets of orthogroup–phenotype matrices, noting that each species pair requires a different gene-to-orthogroup translation. Rows with the same contents, which might be thought of as representing in-paralogous phenotypes, are collapsed together; consequently, all rows in a matrix are unique.

Within the resulting matrix, an all-versus-all search is performed, identifying all pairs of phenotypes which overlap. The selected operation (*and* or *not*) is carried out upon each overlapping pair. The latter operation is carried out twice, *e.g., A – B* and *B – A.* If a resulting pseudo-phenotype is the same as either of the inputs, or consists of fewer than three orthogroups, it is discarded. Phenotypes with the exact same orthogroup sets are merged, as before.

Lastly, phenologs are identified between the target species and the computed matrix (as in [4]). A filter is imposed, throwing away all phenologs with an intersection size of less than two. In addition, phenologs which predict no new genes (the intersection consists of the entire Boolean phenotype) are discarded. The orthogroups within the pseudo-phenotype but outside the overlap are translated into target-species gene identifiers and are considered to be candidates.

### 2.2 Correction for multiple hypothesis testing

To address the problem of multiple hypothesis testing, we performed an empirical permutation test similar to that used in [4]. We experimented with two different schemes and found that they produced similar results.

In both schemes, the psuedo-phenotype–orthogroup matrices from Figure 1 are used. In one, a single permutation is generated of an entire matrix and each row (phenotype) has its contents rearranged according to that same permutation. In the other scheme, each row is permuted separately.

**Figure 1:**
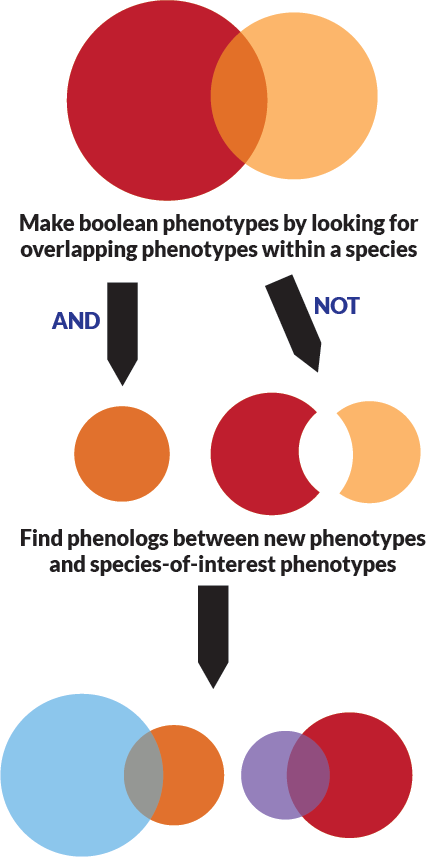
Process for the identification of Boolean phenologs. While Boolean combinations of phenotypes may be identi_ed using any of the Boolean operators, we examine and and not combinations, as these are likely to provide relatively narrower views of the underlying conserved processes.

In each scheme, phenologs are calculated for the matrix permutation; the process is repeated 1,000 times. The resulting distributions are plotted as in Figures 2-3, and a positive predictive value is calculated [9] for every phenolog as in [4].

**Figure 2:**
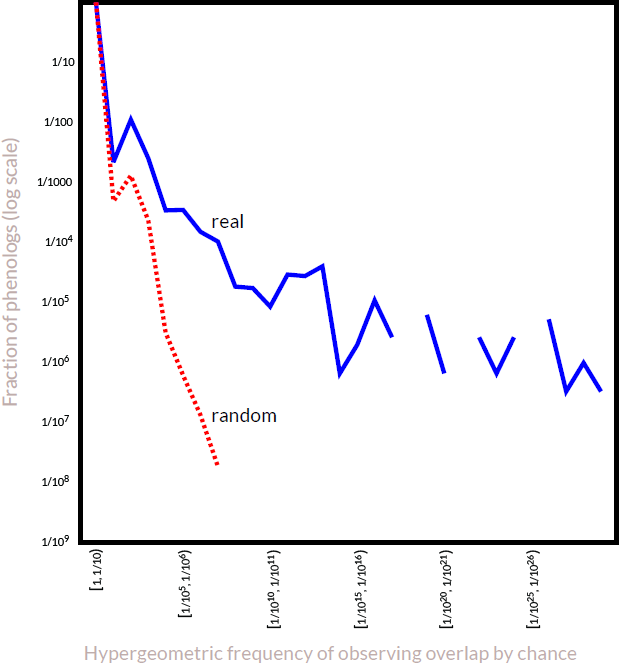
Real and null distributions based on a permutation test of Human–Zebrafish Boolean not phenologs. The solid blue line represents the real distribution of *p* values between phenotypes in a target species and Boolean phenotypes in a source species. The null distribution is based on 1,000 independent runs, permuting each matrix as a unit rather than each row independently. The distributions shown are for predicting human from Zebrafish using the *not* operation.

**Figure 3:**
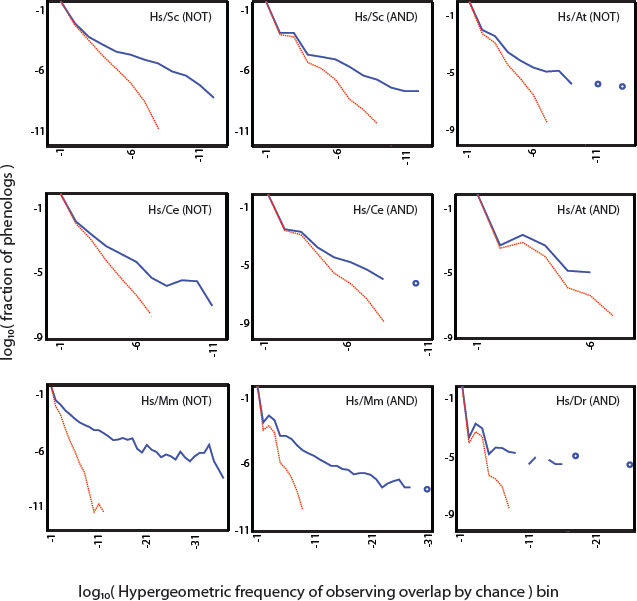
Real and null distributions based on a permutation tests of Boolean phenologs. As with the human–zebrafish not permutation test, the real data offer more extreme hypergeometric frequencies than randomized data, with a generally greater separation between real and random on the *not* combinations than set intersection phenologs. Each x axis bin spans a single log_10_ order of magnitude, with the labels corresponding to the more negative exponent in the bin boundary. The solid blue lines and circles represent the real distribution of *p* values between phenotypes in a target species and Boolean phenotypes in a source species. The null distributions, the dashed red line, are based on 1,000 independent runs, permuting each matrix as a unit rather than each row independently. The distributions shown are for predicting human (Hs) from mouse (Mm), yeast (Sc), *A. thaliana* (At), *C. elegans* (Ce), and zebrafish (Dr), using the *not* and *and* operations.

Since the results were similar, we elected to use the single-permutation scheme, as it preserves the joint distribution of genes and phenotypes.

## 3 Results

### 3.1 Hypothesis

Diseases may consist of multiple phenotypes; such diseases are called syndromes. The definition of “phenotype” requires that they be observable [10], but we wondered if it might be helpful to separate phenotypes — as syndromes may be divided — into components. The components might be thought of as potentially arising from errors in distinct functional modules of genes. We hypothesized that it might be possible to identify such modules by performing various mathematical set operations (*and, or, xor, not*) on overlapping phenotype gene sets. Reasoning that *or* (∪, union) and *xor* would produce broader rather than narrower views of phenotype orthology, we elected to consider *and* (∩, intersection) and *not* (−, subtraction) most carefully (Figure 1). In this paper, we present tests of the Boolean phenolog computational strategy and describe several examples in detail, accompanied by literature support for the predicted candidate genes where available.

### 3.2 Increased brown adipose tissue from *Arabidopsis*

To demonstrate the power of this method, we present the first highly ranked phenotype prediction from the first test run (mouse phenotypes from *Arabidopsis* combination phenotypes using the Boolean *and* operation). The mouse phenotype *increased brown adipose tissue amount* is well-predicted from the intersection of the plant GO biological processes *response to salt stress* ∩ *response to sucrose stimulus* (Figure 4).

**Figure 4:**
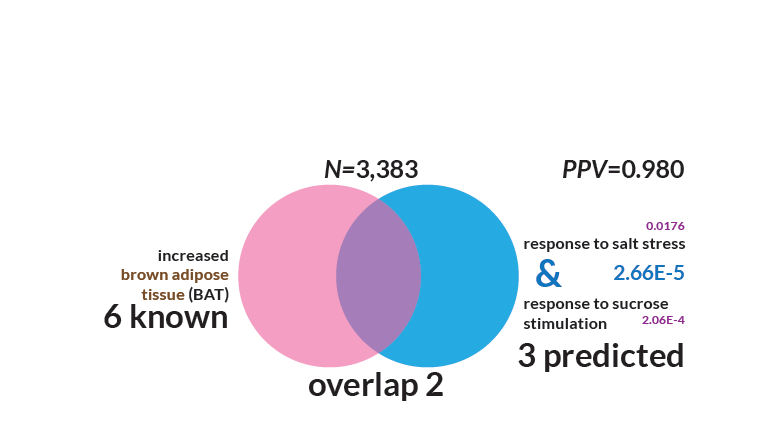
Mouse increased brown adipose tissue genes may be predicted by Arabidopsis Boolean phenotype response to salt stress∩ response to sucrose stimulus. This figure represents the first mouse result outputted by the search of *Arabidopsis* phenotype intersections. The given intersection is the single nearest neighbor. The probability of seeing such an overlap (or larger) by chance is *p* ≤ 10^−5^ (FDR = 0.980), smaller by several orders of magnitude than the probability of seeing either of the individual phenologs separately (*p* ≤ 10^−2^ for salt stress alone and 2 x 10^−4^ for sucrose stimulus, neither of which meets our significance threshold of *p* < 10^−4^). The intersection size is two, with set sizes of six and five; thus, three orthogroups are predicted for the *increased brown adipose tissue* phenotype.

The most consequential difference between brown and white adipose tissue (BAT and WAT) is that the former dissipates energy as heat while the latter stores it. The uncoupling of mitochondrial oxidative phosphorylation in BAT is accomplished by uncoupling protein *UCP1*, the molecular site of non-shivering thermogenesis [11]. Drug-induced uncoupling has been pursued as a treatment for obesity, sometimes with lethal consequences [12,13]. A better understanding of BAT versus WAT physiology might potentially be useful for addressing the obesity epidemic.

The three genes predicted for increased BAT were *Pgd* (6-phosphogluconate dehydrogenase), *Psmd4* (part of the 26S proteasome), and a large orthogroup of homeobox proteins (including *Pitx1−3*, *Isx*, *Pax2-8*, *Rax*, *Alx3*, *Esx1*, *Crx*, *Otx1/2*, *Phox2a/b*, and *Sebox*).

Phosphogluconate dehydrogenase is a lipogenic enzyme in the pentose phosphate pathway, and seems to be expressed in WAT, BAT, and liver, with activity varying according to sex and tissue [14]. Pankiewicz *et al.* also found that oestradiol regulates liver *Pgd* expression; whereas Puerta *et al.* found that oestradiol decreased BAT thermogenesis only in cold-acclimated rats [15]. While the full story would require a full literature review to elucidate, *Pgd* seems to be an interesting candidate.

The case for *Psmd4* is only slightly clearer. UCP1 ubiquitinylation is associated with BAT in cold-acclimated animals, and ubiquitinylation seems to control the rate of UCP1 turnover by the proteasome [16]. It is therefore possible that *Psmd4* plays a direct role in BAT thermogenesis.

Regarding predictions in the third group, homeobox genes are involved in adipogenesis, but it’s unclear whether any of the candidates play a role; these too may be worthy of additional exploration.

In short, candidate genes can be predicted from plant combination phenotypes at high predictive values (PPV = 0.980) and may be worthy of additional exploration.

### 3.3 Set difference: methylmercury and breast cancer

Next, we looked at Boolean combinations consisting of set differences, reasoning that these would identify more narrowly-defined modules rather than the larger assemblies of processes in which those specific modules play a role.

Perhaps the most striking example of the applicability of this technique, and of Boolean phenologs in general, is the case of human breast cancer being predicted from *response to methylmercury* less *response to metal ion* (PPV = 0.992, *p* ≤ 10^−5^; Figure 5). We expected that the genes predicted would by DNA repair-related, as with many cancer phenologs (for example, breast cancer has a phenolog in plants with the intersection of *DNA repair* and *response to gamma radiation, p* ≤ 10^−5^, with two predicted orthogroups: *ATR* and *ERCC6*, which both appear to be DNA repair genes). Instead, these genes highlight two other pathways by which organisms (or individual cells) suppress cancer growth: apoptosis and oxidative stress response.

**Figure 5:**
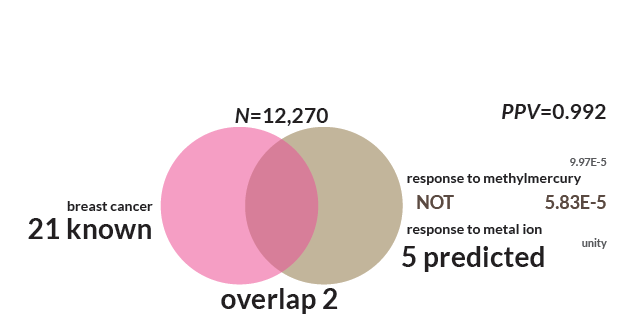
Breast cancer is predicted from genes involved in zebrafish methylmercury response but not response to metal ion. Five genes were predicted for breast cancer. The phenolog overlap was of size two, and nineteen known breast cancer genes were missed. The probability of seeing an overlap between breast cancer and defective methylmercury response alone is *p* < 10^−4^.

Methylmercury (MeHg^+^ or just MeHg) is an organometallic cation, so this example appears at first glance to be a strange combination. However, the *response to methylmercury* GO annotation is a child node of *response to organic substance* in the directed acyclic graph, not *response to metal ion.* A recent article by McElwee *et al.* examines the effects of organic (MeHgCl) and inorganic mercury (HgCl_2_) on *C. elegans,* finding eighteen genes which were important to mercurial exposure response — and only two which responded to both types of mercury [17]. The mechanisms for mercury toxicity are incompletely understood, but it seems clear that even if the two mechanisms are the same or similar (as argued by Clarkson *et al.* [18]), the organismal and cellular responses differ. Thus, we believe this phenolog to be consistent with the findings of McElwee et *al.* [17].

Five genes are predicted: *MT-COI*, *JUN* (c-Jun), *SOD2*, *GADD45B*, and *BAX*. All of these genes — *MT-COI* especially — appear to be involved in the apoptotic response to reactive oxygen species (ROS). The mechanisms by which methylmercury generates ROS are not entirely clear, as previously mentioned, but are reviewed by Farina *et al.* [19]. Generally, MeHg^+^ forms a complex with lower-weight thiol and selenol groups, especially glutathione (reviewed separately in [20]). When such groups occur in mitochondrial creatine kinase or respiratory chain proteins, the compound can inhibit mitochondrial function, leading to depolarization of that organelle’s membrane and overproduction of ROS [19].

In the case of a malfunction of any one of the predicted genes, the effects of ROS may be magnified. Mitochondria rely on *SOD2* (Mn-SOD; reviewed in [21]) for conversion of extremely toxic superoxide into molecular oxygen and hydrogen peroxide, which can be eliminated by catalase. Several of the genes (*JUN*, reviewed in [22]; and *GADD45B* and BAX, reviewed in relation to p53 in [23]) play well-characterized roles in stress-induced apoptosis, triggered in the event the cell is overwhelmed by free radicals (or other agents which cause damage).

*MT-COI,* part of complex IV of the oxidative phosphorylation pathway, is the site of an extremely common germ line mutation in cancer patients, which seems to predispose those individuals toward developing cancer [24]. The occurrence of this mutation may suggest a role for this cytochrome C oxidase component in the pro-apoptotic pathway.

Induction of apoptosis is a key route by which chemotherapy targets cancers — and by which cancers circumvent chemotherapy. The roles of Bcl-2 and Bax are reviewed in [25]. Notably, Bcl-2 binds Bax (the protein products of *BCL-2* and our prediction BAX, respectively), and *BCL-2* over-expression confers chemotherapy resistance. When Bcl-2 is low or absent, however, Bax homodimerizes, leading to cell death. Bax is also thought to interact with the mitochondrial voltage-dependent ion channel, and is directly induced by p53 in response to DNA damage.

Interestingly, a model already exists that might explain these breast cancer gene predictions. Martinez-Outschoorn *et al.* proposed, in 2010, the “autophagic tumor stroma model of cancer.” Essentially, the model hypothesized that tumors induce oxidative stress in the tumor microenvironment in order to cause stromal cells to release nutrients which are used for cancer growth [26,27]. Indeed, Trimmer *et al.* found that loss of Caveolin-1 (Cav-1) — whose stromal presence is a strong predictor of survival — dramatically promotes breast cancer growth. Loss of Cav-1 may be rescued by over-expression of *SOD2,* another tumor suppressor, which relieves oxidative stress by processing mitochondrial superoxide radicals. [27]

*GADD45B* also plays a pro-apoptotic role, and was found to be downregulated in at least two cases of hepatocellular carcinomas (reviewed in [28]). Both *GADD45B* and to a lesser extent *SOD2* were observed to be up-regulated in inflammatory breast cancer (IBC) [29]. The differential regulation argues for a *GADD45B* cancer role, whether as an oncogene or a tumor suppressor; and evidence in mice suggests that all of the Gadd45 genes, including *Gadd45b*, are involved in cancer immune response and are tumor suppressors [30].

These observations suggest further investigations into *MT-COI*’s role in apoptosis and oxidative stress response — and hint that all of the predicted genes may be involved in a tumor suppression transcriptional program.

### 3.4 Myopathy from Arabidopsis

Myopathy is a broad category of disease categorized by muscular weakness with twenty-one associated human genes; three of these genes (*DYSF*, *ACTA1*, and *FHL1*) are members of human–plant orthogroups.

Using the Boolean analysis, we find that the combined *Arabidopsis* phenotype *response to red light* less *response to water deprivation,* predicts the involvement of five orthogroups in myopathy (PPV = 0.991; *p* ≤ 10^−5^; Figure 6). The first of these orthogroups is comprised of members of the SWI/SNF complex and Mediator, including *MED25*, *ARID1A (BAF250A), ARID1B (BAF250B*), and *PTOV1*. *MED25* has been observed in a family with Charcot–Marie–Tooth (CMT) syndrome, including childhood onset distal muscle weakness [31].

**Figure 6:**
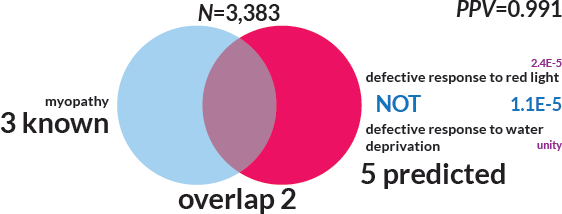
Myopathy is predicted from plant genes involved in red light response but not response to water deprivation. Among 3,383 human-plant orthogroups, three are involved in myopathy; five in the Boolean phenolog *defective response to red light* less *defective response to water deprivation*; and two in the intersection (p ≤ 10^−5^, *PPV* = 0.987). The sub-phenolog myopathy / red light has *p* < 10^−5^.

*ARID1A* and *ARID1B* are both muscle-related. In knockouts of the former, relatively fewer skeletal muscle cells differentiate from embryonic stem cells compared to other differentiation products [32]. *ARID1B* is associated with Coffin-Siris syndrome, which includes hypotonia among its symptoms [33].

In the second orthogroup, two of the three genes are involved in muscle. *CREM* is expressed in ventricular myocytes [34] and *ATF1* is a hypoxia-responsive transcriptional activator of skeletal muscle via mitochondrial *UCP3*[35]; the third, *CREB1,* is expressed in fibroblasts and not in myocytes [34].

The third set of predictions is *TFEB, TFEC, TFE3,* and *MITF*. *TFEC* is a transcriptional activator of non-muscle myosin, and so is ruled out — but is nevertheless an interesting prediction. Over-expression of *TFEB* relieves Pompe disease, a disability of heart and skeletal muscles, by stimulating autophagy [36]. *MITF* has a known autophagy role resembling that of TFEB [37] and is involved in a variety of developmental processes, but no evidence exists suggesting it or *TFE3* are involved in myopathy.

An autophagy link for these predictions is further supported by the fourth predicted orthogroup, consisting of a number of cytochrome P450 (family 2) genes: *CYP2A6–17/13, CYP2B6, CYP2C8–9/18–19*, and *CYP2D6/E1/F1/J2/S1/U1/R1/W1*. Cytochrome P450 is the site of a number of drug interactions — notably with grapefruit, cranberry, and pomegranate juice, which inhibit *CYP3A4,* a metabolizer of statins. *CYP2C8, CYP2C9,* and *CYP2C19* are involved in various statin-induced myopathies [38-40]. At least with the last of these, the mechanism is autophagy related [41]. Similarly, predicted gene *CYP2D6* increased statin efficacy and is a predicted drug interaction site with *3A4* [42]. Finally, *CYP2E1* metabolizes ethanol — which also causes myopathy — and inhibits autophagy [43-45].

Two genes for which we found no literature support are *FAM50A* and *FAM50B,* predicted as a single orthogroup; neither appears to be particularly well researched. These may be good candidates for autophagy genes.

There are at least two other Boolean phenologs of myopathy (both intersectional rather than subtractive) at *p* ≤ 10^−6^ (*PPV* = 0.987). The first of these is *response to light stimulus*∩*response to red light (p* ≤ 10^−4^ and 10^−5^ for the individual components), which predicts the cytochrome P450 orthogroup. The second is *response to auxin stimulus*∩*response to light stimulus* (the former component has *p* ≤ 10^−4^), predicting *GHDC*. We found no literature support for *GHDC,* a gene about which little is known; it may thus be an interesting candidate for myopathy and autophagy.

### 3.5 Holoprosencephaly

Finally, in order to demonstrate predictions from multiple phenologs and to show the utility of intersection phenologs, we present an example of a disease with many phenologs among the *and* (intersection) pseudo-phenotypes.

The concept of *k* nearest neighbors-based ranking discussed in previous work [5] is analogous to the *k* =1 case of Boolean *or* (set union) phenologs. We present here an example of *and* (set intersection) phenologs for *k* = 1, with candidate genes for holoprosencephaly (HPE) predicted from *D. rerio* intersection phenologs (see Table 1). In this case, many phenologs are supported by similarly good *p* values, and predict a variety of genes (see Table 2 for *k*NN-based rankings). It is worth noting that no single component’s *p* value is better than the Boolean *p* value.

**Table 1:**
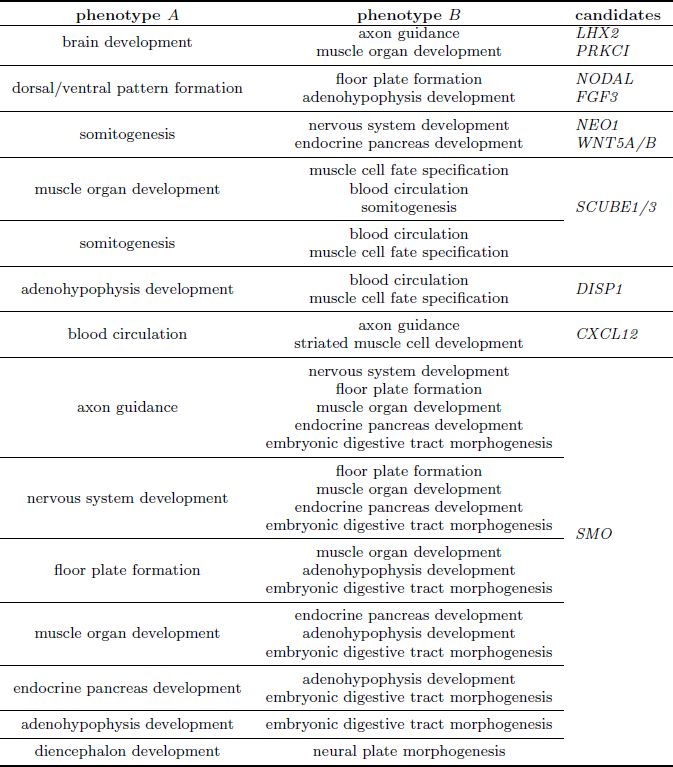
Holoprosencephaly genes are predicted by many intersection phenologs at the same *p* value. We include holoprosencephaly as an example because it has many Boolean *and* phenologs from zebrafish at the same *p* value (6 x 10^−7^, which is the lowest *p* value meeting the filtering criteria). Phenotypes A and B are arbitrarily labeled and indistinguishable overall (but not within individual pairs of table rules). Consider the first entry: brain development *and* axon guidance predicts gene *LHX2*; but brain development *and* muscle organ development predicts *PRKCI.* Nineteen different Boolean phenologs predict *Smoothened.* Rows are grouped first by candidate genes, but also by phenotype A when candidate genes differ. PPV = 0.999.

**Table 2:** Holoprosencephaly candidate genes may be ranked by evidence. We ranked by hand the genes suggested for testing from the phenotypes in Table 1 as they would be ranked by either the naïve Bayes or additive classifiers in the previous Chapter. We included the magnitude (the number of times the gene is predicted by a phenolog in Table 1). Genes within ranks are predicted with the same score (*e.g., NEO1* and *LHX2*) and are thus equally likely under this model). The best predicted gene is *SMO,* also known as *Smoothened.*

| orthogroup members | magnitude |
| --- | --- |
| *SMO* | 19 |
| *SCUBE1/3* | 5 |
| *DISP1* | 2 |
| *CXCL12* | 2 |
| *NEO1* | 1 |
| *WNT5A/B* | 1 |
| *FGF3* | 1 |
| *NODAL* | 1 |
| *LHX2* | 1 |
| *PRKCI* | 1 |

The most highly predicted gene is *SMO* (Smoothened), which is indeed an HPE gene — as determined by Rosenfeld *et al.* in 2010 [46], subsequent to the creation of our database. Rosenfeld *et al.* also identified *DISP1*, ranked third. *NODAL,* one of the bottom-ranked genes, has been observed as promoting an HPE-like phenotype in chick embryos [47] and mice (reviewed in [48]). Another gene ranked with *NODAL, LHX2 (Lim1* /*Lhx2*), is required for mouse head formation [49] and seems to regulate the development of the midline of the brain in that species [50].

The second-ranked orthogroup, consisting of *SCUBE1* and *SCUBE3,* is suggested for HPE candidacy by its role in mouse brain formation (specifically *SCUBE1*) [51], but does not appear to be directly associated with HPE.

### 3.6 Discussion

Boolean phenologs offer a marked improvement over the *k* nearest neighbors approach described previously [5]. The basis for the improvement is not entirely clear, but may be partially subjective: we selected preferentially those Boolean phenologs where the *p* value is less than both the components. Nonetheless, for the *not* phenologs, *all* Boolean *p* values are more significant at our maximum *p* threshold of 1 x 10^−4^. This property is made more likely, but not guaranteed, by the mathematics of set intersection probabilities. If the subtracted phenotype intersects with the query, the Boolean phenotype will have a smaller intersection as a result of the subtraction; and thus the *p* value is more likely to rise above our threshold. The *p* value of two sets with no intersection is within ∈ of unity.

A minor contributor is the way in which the genes in the single-species matrices are translated into orthogroups. For example, plant phenotypes *regulation of telomere maintenance* and *regulation of chromosome organization* have differing sets of genes; but when the genes are translated into human–plant orthogroups, and those plant genes without human orthologs are removed, the two phenotypes collapse into one (these are in-paralogs in *A. thaliana* with respect to *H. sapiens*).

The filtering procedure also plays a role. The original phenolog method looked for any intersection at all; we require an intersection of size two or larger, and additionally that at least one new orthogroup is predicted. For *and* phenotypes, these last two criteria imply that the intersection between the Boolean components must be at least size three.

The probability of finding such a three-way intersection by chance is quite low, and would ordinarily be described by the multivariate hypergeometric distribution (thus, the given probabilities may be conservative over-estimates). The frequency of the three-way intersections — higher than would be expected at random — is a product of the way in which gene–phenotype associations are discovered by biologists.

It follows, then, that the standard hypergeometric distribution may not be conservative enough for phenologs that are “circular”. We define circular by way of an example. Bardet–Biedl syndrome candidate genes which were identified subsequent to the assembly of our gene–phenotype association database are perfectly predicted by the zebrafish Boolean phenotype *melanosome transport* less *embryonic specification defect* (*p* ≤ 10^−25^). However, zebrafish melanosome transport is studied at least in part for its role in Bardet–Biedl syndrome. As such, melanosome transport-associated genes at the time of database construction were already only one patient validation away from being Bardet–Biedl-associated genes. Circularity may explain the unusually extreme hypergeometric *p* values in the human–mouse Boolean phenologs (see Figure 3).

An additional concern is over-training. One might argue, for example, that by subtracting every phenotype from every other phenotype, and then looking for phenologs, one is simply trimming away some of the less immediately relevant portions of phenotype gene sets; however, Boolean phenologs simply provide good hints about which genes and processes should be studied for a deep homology role. Some genes may be missed, but the advantage is an observably lower false positive rate. As with any predictive methodology, we view Boolean phenologs as hypothesis generators, suggesting starting points for deeper investigation.

A final issue is confirmation bias, particularly when pursuing literature validation. Ioannidis argued in 2005 that most published research findings are false positives, in part due to the failure of researchers to share negative results (which are more likely to be correct) [52]. Indeed, our gene–phenotype association matrices only contain “positive” and “unobserved,” and lack negatives. Negatives would be extremely useful for evaluation of results as well. Particularly when predicting extremely well-studied diseases like cancer, it’s unlikely that one will find a well-characterized gene which is not in some way associated if one looks long enough. One potential approach might be is to silently insert an additional random prediction in some set fraction of phenologs selected for literature validation, and then determine how frequently the random prediction is marked as true by the researcher.

## 4 Conclusions

Boolean phenologs offer a computational approach to calculate phenologs [4,5], in a manner designed to focus attention on specific component modules and subprocesses which underlie diseases and phenotypes, as well as the non-obvious homologies which exist between organisms. Here, we have presented Boolean phenolog models for human diseases such as myopathy and breast cancer, as well as increased brown adipose tissue in mice. We describe a number of predictions worthy of further testing, including *FAM50A* and *FAM50B* for autophagy and *MT-COI* for oxidative stress and apoptosis in cancer.

Notably, as with phenologs, the Boolean approach offers insight into those elements of diseases, traits, or processes which are conserved and well-studied. A human–zebrafish phenolog is informative about breast cancer only insofar as cancer is affected by well-characterized processes shared between the two organisms. Phenologs are incapable of highlighting the uniquely human components of breast cancer, but can give us information about the roles of oxidative stress and apoptosis — or about DNA repair genes — in cancer.

## Availability of source code and data

Gene–phenotype associations are available at the phenologs website, phenologs.org. The source code and documentation are available on GitHub at http://github.com/marcottelab/boolean

## Authors’ contributions

J.O.W. performed the analyses and drafted the manuscript. M.T. performed preliminary analyses. E.M.M. supervised the research. All authors edited the final manuscript.

## Acknowledgements

This work was supported by grants from the Cancer Prevention Research Institute of Texas, the National Science Foundation, the National Institutes of Health, and the Welch Foundation (F–1515); and a National Science Foundation Graduate Research Fellowship (to J.O.W.).

